# Identification of biological mechanisms by semantic classifier systems

**DOI:** 10.1101/335737

**Authors:** Ludwig Lausser, Florian Schmid, Lea Siegle, Rolf Hühne, Malte Buchholz, Hans A. Kestler

## Abstract

The interpretability of a classification model is one of its most essential characteristics. It allows for the generation of new hypotheses on the molecular background of a disease. However, it is questionable if more complex molecular regulations can be reconstructed from such limited sets of data. To bridge the gap between complexity and interpretability, we replace the de novo reconstruction of these processes by a hybrid classification approach partially based on existing domain knowledge. Using semantic building blocks that reflect real biological processes these models were able to construct hypotheses on the underlying genetic configuration of the analysed phenotypes. As in the building process, also these hypotheses are composed of high-level biology-based terms. The semantic information we utilise from gene ontology is a vocabulary which comprises the essential processes or components of a biological system. The constructed semantic multi-classifier system consists of expert base classifiers which each select the most suitable term for characterising their assigned problems. Our experiments conducted on datasets of three distinct research fields revealed terms with well-known associations to the analysed context. Furthermore, some of the chosen terms do not seem to be obviously related to the issue and thus lead to new, hypotheses to pursue.

**Author summary:** Data mining strategies are designed for an unbiased de novo analysis of large sample collections and aim at the detection of frequent patterns or relationships. Later on, the gained information can be used to characterise diagnostically relevant classes and for providing hints to the underlying mechanisms which may cause a specific phenotype or disease. However, the practical use of data mining techniques can be restricted by the available resources and might not correctly reconstruct complex relationships such as signalling pathways.

To counteract this, we devised a semantic approach to the issue: a multi-classifier system which incorporates existing biological knowledge and returns interpretable models based on these high-level semantic terms. As a novel feature, these models also allow for qualitative analysis and hypothesis generation on the molecular processes and their relationships leading to different phenotypes or diseases.

## Introduction

Understanding the information hidden in molecular profiles brings new possibilities for the accurate identification of diseases and the selection of individual treatments. It is one foundation of precision or personalised medicine [1]. Molecular profiles typically do not permit any direct inspection in a multivariate setting, as they usually comprise tens of thousands of measurements. Computer-aided systems are paramount for interpretation and categorisation.

Such a computer-aided system is a classification system that assigns known phenotypes, called classes to molecular profiles. The classification system itself generates the diagnostic rules applied in this process. The corresponding decision criteria are constructed in a training phase according to a set of sample instances. Data-driven classification algorithms therefore generate new hypotheses on the underlying characteristics of a biological phenotype and allow for addressing different research questions. Nevertheless, this ability can be limited by the restricted amount of training samples. Complex molecular interactions might not be detected.

However, information about molecular processes can be extracted from various sources such as biological literature and databases [2]. A continuous challenge is to incorporate this heterogenous information into existing classification algorithms. This will lead from purely data-driven systems to knowledge-based modelling approaches. An overview of existing approaches can be found in Porzelius et al. [3]. In the context of feature selection, the knowledge about the construction of biological processes can be used to guide the selection of single features. For example Binder et al. utilise the definition of molecular pathways to guide the construction of a boosting ensemble [4]. Johannes et al. extract information from protein-protein interaction networks for the optimisation criterion of their recursive feature elimination algorithm [5]. Another possible use for knowledge-based feature selection is the en bloc evaluation of the features of a term. Abraham et al. generate an intermediate representation by combining the measurements of a process into a single centroid [6]. Lottaz et al. designed a hierarchical system that is guided by the terms of the gene ontology [7]. Taudien et al. utilise semantic terms for aggregating mutation patterns [8]. In this work, we concentrate on semantic multi classifier systems (S-MCS), which constructs an ensemble of term-dependent base classifiers [9]. We extend this approach to incorporate semantic relationships given by an ontology during the training process. This eventually leads to the identification of new biological mechanisms underlying the discrimination process.

## Materials and methods

Classification is the procedure of categorizing an object into one of *k ≥* 2 distinct classes *𝒴* = *{y*_1_, *…, y* _*k*_ *}*. These classes are assumed to reflect some kind of semantic interpretation (e.g. *carcinoma* vs. *inflammation*). They are predicted according to a vector of measurements **x** = (*x*^(1)^, *…, x*^(*n*)^) ^*T*^ of the object. The corresponding decision rule is described by a classification model

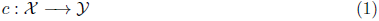

where *𝒳 ⊆* ℝ^*n*^ denotes the measurement or feature space.

The final structure and the properties of a classification model are dependent on the chosen type of concept or function class *𝒞* and the chosen adaptation process *l* to be used on a set of learning instances *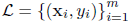*

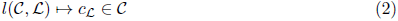

In this basic version, the training procedure *l* can be seen as a purely data-driven process in which a classifier is adapted according to an optimisation criterion. Knowledge about the underlying biology is typically not incorporated.

After the initial training, the generalisation performance is tested in an independent set of test samples 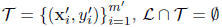. A typical measure of a generalisation ability of a classifier is its accuracy in predicting the class label of an unseen sample correctly:

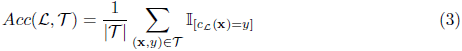

### Interpretability

A second, more fuzzy property of a trained classification model is its ability of providing insight into the characteristics, causes or processes of a difference between classes. We will call this property *interpretability* in the following. Interpretability might be seen as a model’s ability of generating hypotheses on the underlying data.

This property is partially influenced by the structure and complexity of the chosen concept class *𝒞*, which determines the syntactical structure of the final decision rule [10]. For example, a logical conjunction can be more easily interpreted than a linear combination [11].

The interpretability of a classifier is also influenced by the interpretability of its input signals and their transformations [12–14]. A binary signal (e.g. high/low) often allows an easier interpretation than a real-valued gradient [15]. Nevertheless, many of these techniques do not directly provide a possible interpretation regarding the original experiment. While many algorithms lead to a solution with a clear mathematical structure, they often lack a direct semantic interpretation in the terms of the underlying (biological) processes.

### Feature selection

Feature selection is one of the most prominent techniques for increasing the interpretability of high-dimensional profiles [13, 16, 17]. It is typically applied to reduce noisy or irrelevant measurements [18, 19]. By fulfilling the chosen quality criterion the remaining features are considered as possible (informative) candidate markers.

Technically, feature selection can be seen as a process which maps from the chosen type of concept class *𝒞* and the training set *ℒ* to the space *ℐ* of repetition-free index vectors of maximal length *n*

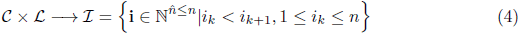

The elements (*signatures*) **i** *∈ ℐ* will be used to indicate the features which pass the selection process **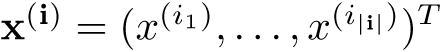**, where **i** = (*i*_1_, *…, i* _*|***i***|*_) ^*T*^. This selection of elements is called a signature. In general, *|ℐ|* = 2 ^*n*^ *-* 1 distinct signatures can be constructed. The ultimately selected signature **i** *∈ ℐ* is typically determined in a *data-driven* or *model-driven* process, which incorporates stochastic or heuristic elements. Exhaustive and therefore optimal strategies are typically avoided due to the size of the corresponding search spaces [20].

Although data- or model-driven approaches can improve the accuracy of a classification procedure the corresponding signatures often lack an overall semantic interpretation. Designed for optimising quality scores, classical feature selection procedures often neglect existing domain knowledge and do not take into account known dependencies or functional relationships among single features. The members of a signature are either selected according to their standalone performance or their contribution to the overall performance of the signature.

An alternative to this approach is the idea of *knowledge-based* feature selection [9]. Here, it is assumed that a predefined set of interpretable signatures *𝒱 ⊆ ℐ* exists. It will be called a *vocabulary 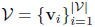* in the following. In this scenario, the problem of constructing a suitable marker signature is altered to a selection problem in which a predefined signature **v** *∈ 𝒱* or a combination of predefined signatures *𝒱* ^*′*^ *⊆ 𝒱* is chosen from a vocabulary

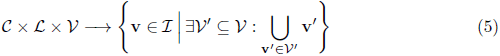

Assuming that *|𝒱| « |ℐ|*, the use of a vocabulary reduces the corresponding search space to 2 ^*|𝒱|*^ *-* 1 elements.

The term “knowledge-based feature selection” refers to the known semantic interpretation of the signatures in *𝒱*. They were designed according to domain knowledge about the measuring process or the subject of an experiment and are assumed to summarise the components of a more complex structure. In general, we will utilise the word *term* to denote both a predefined signature as well as its interpretation. In the context of systems biology, a term might reflect a known molecular signalling pathway [21] or a cellular component [22]. There are pre-formed terms available from ontologies like Gene Ontology (GO) and databases like the Kyoto Encyclopedia of Genes and Genomes (KEGG).

Besides the benefit for the interpretability of individual signatures, the known semantic meaning of the terms also allows the incorporation of domain knowledge about the pairwise relationships between terms. It especially opens the possibility to use semantic networks or ontologies which represent known relationships in form of a graph structure. Similar to the composition of the signatures, these relations must not be learned during a classification experiment. They are typically known a priori and might comprise information which can not be deduced from *ℒ*.

Although we assume that the signatures in a vocabulary are unique

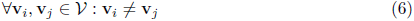

this might not be true for their interpretation. For example, a union of two signatures **v** _*i*_ ∪ **v** _*j*_ might cover the signature of a third one. Therefore we constructed a multi-classifier system that handles the terms and the corresponding measurements individually.

### Semantic multi-classifier system

The design of our semantic multi classifier system (S-MCS) can be seen as an ensemble *E ⊆ 𝒞* _*𝒱*_ of semantic base classifiers *𝒞* _*𝒱*_ which are restricted to the signature of an individual term **v** *∈ 𝒱*

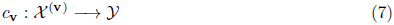

The ensemble itself will be denoted by *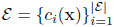*. The restriction to a specific signature will force a classifier *c***_v_**(**x**) to become an expert in interpreting the signals in **v** and therefore to become an expert in interpreting the corresponding term. A S-MCS utilizing a vocabulary *𝒱* will be denoted by S-MCS _*𝒱*_.

The overall structure of a S-MCS is shown in Figure 1d). The individual predictions of the semantic base classifiers are merged on a symbolic level by a late-aggregation strategy [23]

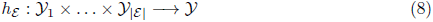

**Fig 1.**
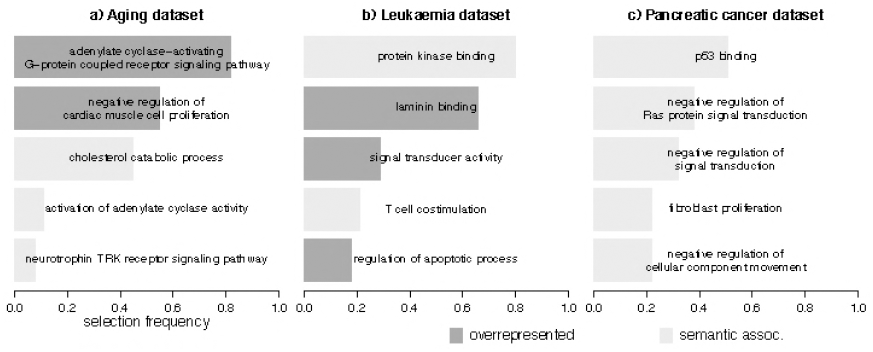
Concept of a semantic multi classifier system (S-MCS): Panels a)-c) show the training steps of a S-MCS _*𝒱*_ *\**. In Panel a) the initial vocabulary *𝒱* is screened for terms that have a significant overlap to the context set **w**. The corresponding sets *𝒱* _*a*_ (**w**) are highlighted in red. They build a set of starting points for an ontological search (Panel b). The direct hyponyms of the terms in *𝒱* _*a*_ (**w**) are utilized as additional candidates for the final selection process (blue). The preselected terms **v** *∈ 𝒱* ^*\**^ are evaluated in a cross validation experiment with semantic base classifiers (Panel c). The top *k* terms with the highest accuracy are chosen for the final classifier ensemble *E*. Panel d) shows the structure of a trained S-MCS. The semantic base classifiers *c*(**x**) *∈ E* are combined via a fusion architecture *h* _*E*_ (**x**).

In this way, the selected signatures are encapsulated during the system’s training and prediction phases. The interpretation of the corresponding terms will not be mixed up. More precisely, our ensemble decision utilises a majority vote on the chosen base classifiers

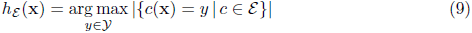

During its training phase the S-MCS will gain access to a preselected set of possible candidate terms *𝒱* ^*\**^ *⊆ 𝒱*. From this set *𝒱* ^*\**^ the terms and the corresponding base classifiers are selected in a data-driven way (Figure 1c). They are chosen iteratively according to their accuracy in a *r × f* cross-validation (CV) on the training set *ℒ*. On the level of terms, this can be formalised as

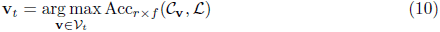

where *𝒱* _*t*_ = *𝒱* _*t-*1\_ *{***v** _*t-*1_ *}* and *𝒱*_1_ = *𝒱* ^*\**^. The initial set of candidate terms *𝒱* ^*\**^ is constructed according to filter criteria based on the available domain knowledge.

### Knowledge-based term selection

A general vocabulary *𝒱* is typically not designed for a particular classification task. It consists of a large collection of widespread terms which can be used to describe several tasks in a certain domain or field. In order to guide the search process of the S-MCS a subset of suitable terms *𝒱* ^*\**^ *⊆ 𝒱* can be pre-selected according to existing domain knowledge.

In the following, we present a term selection strategy which is guided by knowledge about suitable measurements. We assume that an additional signature **w** */∈ 𝒱* exists which comprises known signals that are associated with a given classification task. The pre-selection of *𝒱* ^*\**^ consists of two subsequent steps. Both only utilise part of the information contained in the information contained in the measurements of *𝒯* and the chosen model type *𝒞*.

The first one, an *overrepresentation analysis*, operates directly on the level of feature indices. It neither utilises the values measured in the experiment nor the prior knowledge about the semantic relationships of the terms. It only considers the information about the overlaps of the signatures.

The second one, an *ontological search*, is guided by external domain knowledge. It directly utilises the connections of the corresponding graph structure.

#### Overrepresentation analysis

The overrepresentation analysis is based on a set level statistic which determines the intersect **v** ∩ **w** of two given sets **v**, **w** *⊆ ℐ*. The significance, according to a significance level α, of the overlap between these two sets is determined via Fisher’s exact test [24] on the basis of a hypergeometric distribution

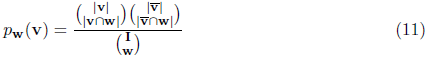

In the context of our S-MCS we utilise overrepresentation analysis as a technique for integrating vocabularies or ontologies (Figure 1a). A term is added to the set of candidates if it has a significant overlap to the initial gene set **w**

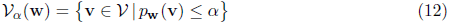

#### Ontological Search

An ontological search relies on the graphical representation of the relationships between the terms of a vocabulary *𝒱*. It requires the availability of an ontology or a semantic network. The structure of the corresponding graph induces a neighbourhood which can be screened for suitable candidate terms later on.

In the following, we concentrate on ontological graph structures (Figure 1b). Thus, we assume a directed hypernym-hyponym relationship *r*(**w**, **v**) which reflects the hierarchical relationship between a category **w** and its subcategories **v**. For a given set of already selected terms *W ⊆ 𝒱* we additionally consider those terms as candidates which are direct subcategories of the terms in *𝒱*

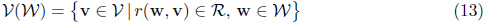

Our final candidate set for the data-driven term selection *𝒱* ^*\**^ is then constructed from the terms selected by the overrepresentation analysis and the ontological search

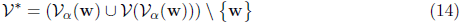

## Experiments

We have evaluated our S-MCS in three molecular settings. A short characterisation of the datasets can be found in Table 1. They are publicly available and can be downloaded from the Gene Expression Omnibus (GEO) [25, 26]. As a preprocessing step a RMA normalisation was applied to all datasets [27]. The one nearest neighbour classifier (1-NN) was chosen as a base classifier for the S-MCS [28] and the ensemble size was set to three base classifiers. For each of the datasets the mean accuracy of the classifiers was estimated in a 10 *×* 10 cross validation (CV) setting [29]. The experiments were conducted with help of the TunePareto Framework [30].

The described classifier system was compared to four types of data-driven reference classifiers. The nearest centroid classifier (NCC), the one nearest neighbour classifier [28] (1-NN), the linear support vector machine [10] (SVM) and the random forest [31] (100 trees, RF-100) were chosen. Each of these classifiers was applied as a standalone classifier on all features as well as in combination with a data-driven feature selection. In the second case, a correlation filter was applied to the classifiers training set *ℒ* to preselect the 100 features with the highest absolute Spearman rank correlation to the class labels.

The experiments also comprise other knowledge-based approaches besides S-MCS _*𝒱*_ *\**(see below). The proposed system was compared to a S-MCS without access to the relationships of the terms in *𝒱* (S-MCS _*𝒱*_). Here, the S-MCS directly selects the top three terms of *𝒱* according to their classification accuracy (Equation 10). A second

**Fig 2.**
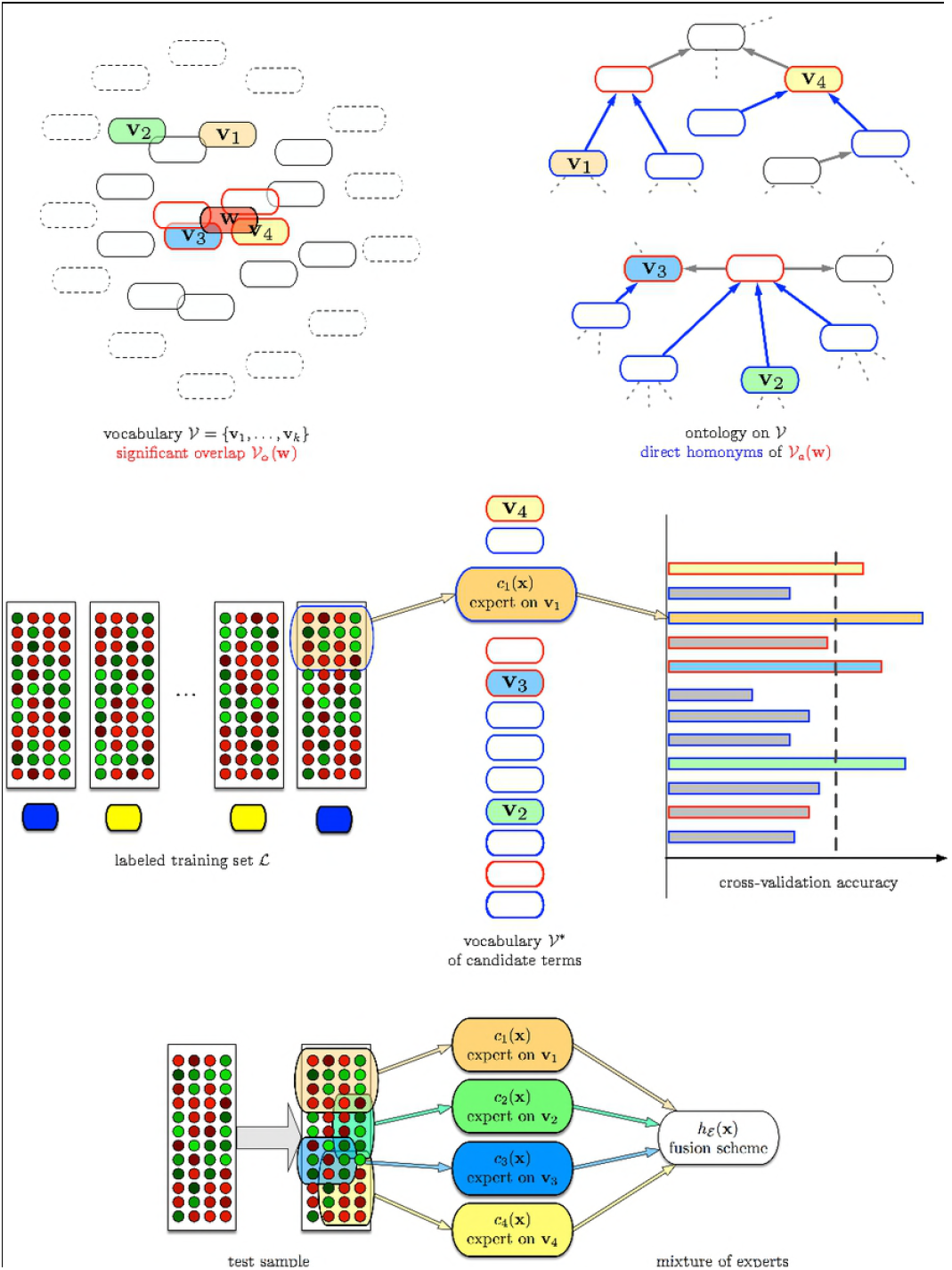
Barplots of the five most frequently selected terms of the S-MCS _*𝒱*_ *\** over all training runs of the 10*×* 10 cross validation. Dark grey bars show terms that have been found via overrepresentation, light grey bars show terms found via ontological search.

S-MCS (S-MCS _*𝒱*_ *α*) is only allowed to access the terms in *𝒱*, which have a significant overlap to the initial seeding set **w** (Equation 12, *α* = 0.05). The proposed system itself will be denoted by S-MCS _*𝒱\**_. For all three S-MCS, the vocabularies will be constructed from the Gene Ontology. Terms are selected according to their accuracy in 3 *×* 3 C𝒱 experiments when training on the training set.

**Table 1.**
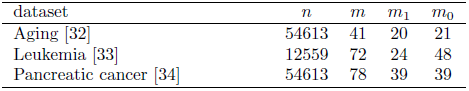
Overview of analysed datasets. The overall number of features *n*, the overall number of samples *m* and the class wise number of samples *m*_1_ and *m*_0_ are reported.

### Gene Ontology

The Gene Ontology (GO) provides three curated vocabularies representing gene product properties [35]: Cellular Components, Molecular Functions, and Biological Processes. Each of these vocabularies is organised in form of a hierarchical ontology. Cellular Component terms describe parts of a cell or its extracellular environment like the nucleus or protein complexes, but not individual proteins or multicellular anatomical structures. Molecular Function terms describe activities of a gene product at a molecular level, for example catalytic activity. They should be unambiguous and signify the same thing in any species. Biological Process terms describe series of events or molecular functions like signal transduction which have at least two distinct steps and a defined beginning and end.

## Results & discussion

The results will be discussed for each dataset individually. An overview of the achieved classification results can be found in Table 2. The most frequently selected terms are given in Fig 2.

### Ageing dataset

Lu et al. provided gene expression data of post-mortem human brain samples of different ages as a part of a study on stress resistance in ageing and Alzheimer’s disease [32]. They divided the samples into four age classes: less than 40 years (12 samples), 40-69 years (9 samples), 70-94 years (16 samples), 95-106 years (4 samples). The dataset is publicly available from the GEO database under accession GSE53890. Total RNA was isolated from cortical grey matter samples from the frontal pole of post-mortem human brains. It was hybridised to Affymetrix U133 plus 2.0 arrays.

For our analysis of ageing we combined the first two classes into a “adult” class (21 samples) and the latter two into an “aged” class (20 samples). The GO term ageing (GO:0007568) has been chosen as initial set **w** for the term filtering procedure. The corresponding set of measurements consists of 114 genes. These are associated with resistance to disease, homeostasis, and fertility.

**Table 2.**
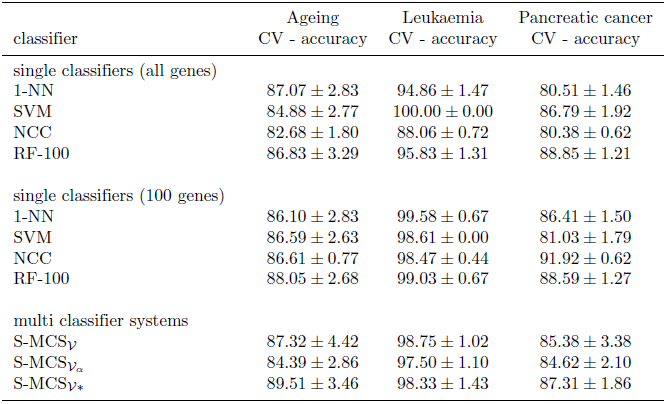
Accuracies of the 10 *×* 10 CV (mn *±* std) for all datasets. Abbreviations: CV, cross validation; mn, mean; std, standard deviation; 1-NN, 1-nearest neighbour; SVM, support vector machine; NCC, nearest centroid classifier; RF-100, random forest with 100 trees; S-MCS, semantic multi classifier system.

| classifier | Ageing<br>CV - accuracy | Leukaemia<br>CV - accuracy | Pancreatic cancer<br>CV - accuracy |
| --- | --- | --- | --- |
| single classifiers (all genes) |  |  |  |
| 1-NN | $87.07 \pm 2.83$ | $94.86 \pm 1.47$ | $80.51 \pm 1.46$ |
| SVM | $84.88 \pm 2.77$ | $100.00 \pm 0.00$ | $86.79 \pm 1.92$ |
| NCC | $82.68 \pm 1.80$ | $88.06 \pm 0.72$ | $80.38 \pm 0.62$ |
| RF-100 | $86.83 \pm 3.29$ | $95.83 \pm 1.31$ | $88.85 \pm 1.21$ |
| single classifiers (100 genes) |  |  |  |
| 1-NN | $86.10 \pm 2.83$ | $99.58 \pm 0.67$ | $86.41 \pm 1.50$ |
| SVM | $86.59 \pm 2.63$ | $98.61 \pm 0.00$ | $81.03 \pm 1.79$ |
| NCC | $86.61 \pm 0.77$ | $98.47 \pm 0.44$ | $91.92 \pm 0.62$ |
| RF-100 | $88.05 \pm 2.68$ | $99.03 \pm 0.67$ | $88.59 \pm 1.27$ |
| multi classifier systems |  |  |  |
| S-MCS<sub>V</sub> | $87.32 \pm 4.42$ | $98.75 \pm 1.02$ | $85.38 \pm 3.38$ |
| S-MCS<sub>V<sub>α</sub></sub> | $84.39 \pm 2.86$ | $97.50 \pm 1.10$ | $84.62 \pm 2.10$ |
| S-MCS<sub>V*</sub> | $89.51 \pm 3.46$ | $98.33 \pm 1.43$ | $87.31 \pm 1.86$ |

Table 2 shows the results of the 10*×* 10 cross validation for the ageing dataset. Operating on the full expression profiles, the standalone classifiers achieve accuracies from 82.68 *±* 1.80% (NCC) to 87.07 *±* 2.83% (1-NN). If the classifiers are coupled to the correlation filter their accuracies vary from 85.61 *±*0.77% (NCC) to 88.05 ± 2.68% (RF-100). The semantic multi classifier systems achieve accuracies in the range of 84.39 *±* 2.86% (S-MCS _*𝒱*_ *α*) to 89.51 *±* 3.46% (S-MCS _*𝒱\**_). Here, the performance of a classifier system using all terms of the ontology (87.32 *±* 4.42%, S-MCS _*𝒱*_) is decreased by a standalone overrepresentation filter but increased by the pipeline of both term filters.

Fig 2a) gives an overview on the 10 *×* 10 cross validation experiments with the S-MCS _*𝒱\**_. The five most frequently selected terms are shown. Two of these terms are related to adenylate cyclase. The term *adenylate cyclase activating G-protein coupled receptor signalling pathway* is the most frequently selected one (82.0%; 42 associated genes). The term *activation of adenylate cyclase activity* was chosen in 11.0% of all experiments (30 genes). The other frequently selected terms are the *negative regulation of cardiac muscle cell proliferation* (55.0%; 7 genes), the *cholesterol catabolic process* (45.0%; 12 genes) and the less frequent *neurotrophin TRK receptor signaling pathway* (8.0%; 275 genes).

As expected due to the common relation to adenylate cyclase the first two terms have an overlap (9 genes). Additionally terms one and five (9 genes) and four and five (8 genes) have an overlap. However, these overlaps are only pairwise. This indicates that the classification performance is not related to only a small subset of genes associated to the terms.

For most of these terms evidence for a connection to ageing can be found in literature. The influence of adenylate cyclase and the activation of G-proteins on the process of ageing is for example discussed by Feldmann et al. [36]. G-proteins are also known for their role in cardiac muscle cell proliferation during aging [37]. This might explain the high selection rate of the negative regulation of cardiac muscle cell proliferation term. Changes in the cholesterol catabolic process are discussed in hippocampal and cognitive ageing across lifespan [38]. Costantini et al. describe the way ageing affects the signalling pathway which regulates TRK receptor signalling [39].

### Leukaemia dataset

The data from Armstrong and co-workers consist of gene expression data of 24 human acute lymphoblastic leukaemia (ALL) and 48 acute myeloid leukaemia (AML) and mixed-lineage leukaemia (MLL) patients [33]. Total RNA was isolated from mononuclear peripheral blood or bone marrow cells of patients at diagnosis or relapse. It was hybridised to Affymetrix U95A arrays. Data is available from the Broad Institute at http://portals.broadinstitute.org/cgi-bin/cancer/publications/pub_paper.cgi?mode=view&paper_id=63.

For the leukaemia dataset an initial signature **w** was extracted from the Kyoto Encyclopedia of Genes and Genomes (KEGG). The union (98 genes) of the descriptions for *chronic myeloid leukaemia* (73 genes) and *acute myeloid leukaemia* (60 genes) was chosen. The overlap of the signatures consists of 35 genes. Both sets contain elements related to PI3K-Akt and MAPK signalling as well as genes involved in cell cycle.

For this dataset the accuracies of the 10 *×* 10 cross validation experiments are listed in Table 2. The standalone classifiers achieve accuracies between 88.06 *±* 0.72% (NCC) and 100.00 *±* 0.00% (SVM). The application of the correlation filter leads to accuracies in the interval between 98.47 *±* 0.44% (NCC) and 99.58 *±* 0.67% (1-NN). The accuracies of the semantic multi classifier systems lie in the range of 97.50 *±* 1.10 (S-MCS _*𝒱*_ *a*) and 98.75 *±* 1.02 (S-MCS _*𝒱*_). For this dataset the application of the filter pipeline slightly decreases the performance (98.33 *±* 1.43%, S-MCS _*𝒱\**_).

The most frequently selected terms of the S-MCS _*𝒱\**_ are given in Fig 2b). In 80% of all experiments the term *protein kinase binding* was chosen (332 genes), followed by the term *laminin binding* (66.0%; 24 genes) and less frequently by the terms *signal transduction activity* (29.0%; 226 genes) and *T cell costimulation* (21.0%; 72 genes). In 18.0% the term *regulation of apoptotic process* was chosen (155 genes).

For this dataset more overlaps of single genes occur than in the others, especially between the most frequently selected term *protein kinase binding* and the other terms. The largest overlaps are between the most frequently selected term *protein kinase binding* and *signal transduction activity* (19 genes overlap), *T cell costimulation* (13 genes overlap), and *regulation of apoptotic process* (10 genes overlap). The first two terms contain a high number of genes. This might result in the especially high overlap between the two terms. The overlaps between the other terms are smaller than 10 genes. All sets (except the first one) contain the LGALS1 gene which is hypermethylated in MLL patients [40] and might be an important feature for the classification problem.

For three of the chosen terms, an association to the subject of leukaemia was shown in literature. Examples for the influence of *protein kinase binding* are given by Vijapurkar et al. [41]. The role of laminin receptors is characterised by Federico et al. [42]. A discussion of signal transduction processes in in the context of leukaemia is given by Ihle et al. [43].

### Pancreatic cancer dataset

Badea et al. [34] analysed gene expression data of 36 pancreatic ductal adenocarcinoma (PDAC) tumour patients. They used pairs of tumour samples and normal pancreatic tissue samples from the same patients. The dataset is publicly available in the GEO database under accession GSE15471. Total RNA was hybridised to Affymetrix U133 plus 2.0 arrays. As an initial marker set a mRNA signature developed from the data published by Gress and co-workers was chosen [44].

Table 2 shows the accuracies achieved on the Pancreatic cancer dataset. Utilising all available features the standalone classifiers achieve accuracies in the range from *±* 0.62% (1-NN) to 88.85 *±* 1.21% (RF-100). Coupled to the correlation based filter the results lie between 81.02 *±* 1.79% (SVM) and 91.92 *±* 0.62% (NCC). The semantic multi classifier systems gain accuracies from 84.62 *±* 2.10 (S-MCS _*𝒱*_ *a*) to 87.31 *±* 1.86% (S-MCS _*𝒱\**_).

The terms most frequently selected by S-MCS _*𝒱\**_ are shown in Fig 2c). The term *p53 binding* was selected in 51.0% of all experiments (56 genes). Base classifiers for *negative regulation of Ras protein signal transduction* (24 genes) and *negative regulation of signal transduction* (18 genes) were chosen in 38.0% and 32.0% of all experiments. The terms *fibroblast proliferation* (4 genes) and *negative regulation of cellular component movement* (6 genes) were both selected in 22.0% of all experiments. The terms selected by the classifier system do not overlap.

The role of p53 in tumor development is frequently discussed in literature [45, 46]. The influence of Ras mutations is for example discussed by Fensterer et al. [47]. Fibroblasts and fibroblast proliferation are also known to play a role in cancer [48, 49]. In case of pancreatic cancer they play a role in the invasiveness of the cancer [50]. This could also be related to the last term which is negative regulation of cellular component movement. Effects on the associated components may increase cell movement which is a key step in the formation of metastases [51–54].

## Conclusion

In this work, we proposed an ontologically guided multi-classifier system that can incorporate semantic domain knowledge into a data-driven modelling process. Utilizing information about the integral parts of a biological process or a cellular component the system was able to construct interpretable hypotheses on the molecular background of the analysed phenotypes or diseases in the form of a set of GO terms. These terms are a high-level description of the problem and a basis for further research.

In our scenarios, we evaluated the multi-classifier system in three well-structured research fields. Literature could corroborate evidence for the chosen semantic terms while also suggesting so far unknown mechanisms. Utilising information about the integral parts of a biological process or a cellular component the system was able to construct interpretable hypotheses on the molecular background of the analysed phenotypes or diseases. Suggesting that the system can be used to guide molecular experiments. One of the central assumptions of systems biology is that common high-level components influence almost all biological processes. Thus, our multi-classifier system can be applied as a general purpose instrument to many different problems in a broad field of applications.

## Acknowledgments

The research leading to these results has received funding from the European Community’s Seventh Framework Programme (FP7/2007-2013) under grant agreement n ^*?*^ 602783, the German Research Foundation (DFG, SFB 1074 project Z1), the Federal Ministry of Education and Research (BMBF, e:Med, SYMBOL-HF, id 01ZX1407A, CONFIRM, id 01ZX1708C) and the Deutsche Krebshilfe (id 111318).

## References

1. Kraus JM, Lausser L, Kuhn P, Jobst F, Bock M, Halanke C, et al. Big data and precision medicine: challenges and strategies with healthcare data. International Journal of Data Science and Analytics. 2018;doi:10.1007/s41060-018-0095-0.

2. Galperin MY, Rigden DJ, Fernández-Suarez XM. The 2015 *Nucleic Acids Research* Database Issue and Molecular Biology Database Collection. Nucleic Acids Research. 2015;43(Database-Issue):1–5.

3. Porzelius C, Johannes M, Binder H, Beißbarth T. Leveraging external knowledge on molecular interactions in classification methods for risk prediction of patients. Biometrical Journal. 2011;53(2):190–201.

4. Binder H, Schumacher M. Incorporating pathway information into boosting estimation of high-dimensional risk prediction models. BMC Bioinformatics. 2009;10:18–28.

5. Johannes M, Brase JC, Fr¨ohlich H, Gade S, Gehrmann MC, Fälth M, et al. Integration of pathway knowledge into a reweighted recursive feature elimination approach for risk stratification of cancer patients. Bioinformatics. 2010;26(17):2136–2144.

6. Abraham G, Kowalczyk A, Loi S, Haviv I, Zobel J. Prediction of breast cancer prognosis using gene set statistics provides signature stability and biological context. BMC Bioinformatics. 2010;11(1):277–291.

7. Lottaz C, Spang R. Molecular decomposition of complex clinical phenotypes using biologically structured analysis of microarray data. Bioinformatics. 2005;21(9):1971–1978.

8. Taudien S, Lausser L, Giamarellos-Bourboulis EJ, Sponholz C, F S, Felder M, et al. Genetic Factors of the Disease Course After Sepsis: Rare Deleterious Variants Are Predictive. EBioMedicine. 2016;12:227–238.

9. Lausser L, Schmid F, Platzer M, Sillanpää MJ, Kestler HA. Semantic multi-classifier systems for the analysis of gene expression profiles. Archives of Data Science Series A (Online First). 2014;1(1).

10. Vapnik V. Statistical Learning Theory. Wiley; 1998.

11. Kestler HA, Lausser L, Lindner W, Palm G. On the fusion of threshold classifiers for categorization and dimensionality reduction. Computational Statistics. 2011;26(2):321–340.

12. Jolliffe IT. Principal Component Analysis. Springer; 2002.

13. Guyon I, Gunn S, Nikravesh M, Zadeh LA. Feature Extraction: Foundations and Applications (Studies in Fuzziness and Soft Computing). Springer; 2006.

14. Lausser L, Schmid F, Schirra LR, Wilhelm AFX, Kestler HA. Rank-based classifiers for extremely high-dimensional gene expression data. Advances in Data Analysis and Classification. 2016;doi:10.1007/s11634-016-0277-3.

15. Müssel C, Schmid F, Blätte TJ, Hopfensitz M, Lausser L, Kestler HA. BiTrinA - multiscale binarization and trinarization with quality analysis. Bioinformatics. 2016;32(3):465–468.

16. Liu H, Motoda H, editors. Computational methods of feature selection. Chapman & Hall/CRC data mining and knowledge discovery series. Chapman & Hall/CRC; 2008.

17. Lausser L, Szekely R, Schirra LR, Kestler HA. The Influence of Multi-class Feature Selection on the Prediction of Diagnostic Phenotypes. Neural Processing Letters. 2017;doi:10.1007/s11063-017-9706-3.

18. Blum A, Langley P. Selection of Relevant Features and Examples in Machine Learning. Artificial Intelligence. 1997;97(1-2):245–271.

19. Guyon I, Elisseeff A. An introduction to variable and feature selection. Journal of Machine Learning Research. 2003;3:1157–1182.

20. Kraus JM, Lausser L, Kestler HA. Exhaustive k-nearest-neighbour subspace clustering. Journal of Statistical Computation and Simulation. 2015;85(1):30–46.

21. Kanehisa M, Goto S. KEGG: Kyoto Encyclopedia of Genes and Genomes. Nucleic Acids Research. 2000;28(1):27–30.

22. Subramanian A, Tamayo P, Mootha VK, Mukherjee S, Ebert BL, Gillette MA, et al. Gene set enrichment analysis: A knowledge-based approach for interpreting genome-wide expression profiles. Proceedings of the National Academy of Sciences of the United States of America. 2005;102(43):15545–15550.

23. Sammut C, Webb G. Encyclopedia of Machine Learning. Springer, New York; 2010.

24. Fisher RA. On the Interpretation of χ^2^ from Contingency Tables, and the Calculation of P. Journal of the Royal Statistical Society. 1922;85(1):pp. 87–94.

25. Edgar R, Domrachev M, Lash AE. Gene Expression Omnibus: NCBI gene expression and hybridization array data repository. Nucleic Acids Research. 2002;30(1):207–210.

26. Barrett T, Wilhite SE, Ledoux P, Evangelista C, Kim IF, Tomashevsky M, et al. NCBI GEO: archive for functional genomics data sets-update. Nucleic Acids Research. 2013;41(D1):D991–D995.

27. Irizarry RA, Hobbs B, Collin F, Beazer-Barclay YD, Antonellis KJ, Scherf U, et al. Exploration, normalization, and summaries of high density oligonucleotide array probe level data. Biostatistics. 2003;4(2):249–264.

28. Fix E, Hodges, Jr JL. Discriminatory Analysis: Nonparametric Discrimination: Consistency Properties. USAF School of Aviation Medicine, Randolf Field, Texas; 1951. Project 21-49-004, Report Number 4.

29. Japkowicz N, Shah M. Evaluating Learning Algorithms: A Classification Perspective. Cambridge University Press; 2011.

30. Mu¨ssel C, Lausser L, Maucher M, Kestler HA. Multi-Objective Parameter Selection for Classifiers. Journal of Statistical Software. 2012;46(5):1–27.

31. Breiman L. Random Forests. Machine Learning. 2001;45(1):5–32.

32. Lu T, Aron L, Zullo J, Pan Y, Kim H, Chen Y, et al. REST and stress resistance in ageing and Alzheimer’s disease. Nature. 2014;507(7493):448–454.

33. Armstrong SA, Staunton JE, Silverman LB, Pieters R, den Boer ML, Minden MD, et al. MLL translocations specify a distinct gene expression profile that distinguishes a unique leukemia. Nature Genetics. 2002;30(1):41–47.

34. Badea L, Herlea V, Dima SO, Dumitrascu T, Popescu I. Combined gene expression analysis of whole-tissue and microdissected pancreatic ductal adenocarcinoma identifies genes specifically overexpressed in tumor epithelia. Hepatogastroenterology. 2008;55(88):2016–2027.

35. Ashburner M, Ball CA, Blake JA, Botstein D, Butler H, Cherry JM, et al. Gene Ontology: Tool for the unification of biology. Nature Genetics. 2000;25(1):25–29.

36. Feldman RD, Tan CM, Chorazyczewski J. G Protein Alterations in Hypertension and Aging. Hypertension. 1995;26(5):725–732.

37. Alemany R, Perona JS, Sánchez-Dominguez JM, Montero E, Cañizares J, Bressani R, et al. G protein-coupled receptor systems and their lipid environment in health disorders during aging. Biochimica et Biophysica Acta (BBA) - Biomembranes. 2007;1768(4):964 – 975.

38. Kadish I, Thibault O, Blalock EM, Chen KC, Gant JC, Porter NM, et al. Hippocampal and Cognitive Aging across the Lifespan: A Bioenergetic Shift Precedes and Increased Cholesterol Trafficking Parallels Memory Impairment. The Journal of Neuroscience. 2009;29(6):1805–1816.

39. Costantini C, Scrable H, Puglielli L. An aging pathway controls the TrkA to p75NTR receptor switch and amyloid ß-peptide generation. The EMBO Journal. 2006;25(9):1997–2006.

40. Juszczynski P, Rodig SJ, Ouyang J, O’Donnell E, Takeyama K, Mlynarski W, et al. MLL-Rearranged B Lymphoblastic Leukemias Selectively Express the Immunoregulatory Carbohydrate-Binding Protein Galectin-1. Clinical Cancer Research. 2010;16(7):2122–2130.

41. Vijapurkar U, Fischbach N, Shen W, Brandts C, Stokoe D, Lawrence HJ, et al. Protein Kinase C-Mediated Phosphorylation of the Leukemia-Associated HOXA9 Protein Impairs Its DNA Binding Ability and Induces Myeloid Differentiation. Molecular and Cellular Biology. 2004;24(9):3827–3837.

42. Federico MHH, Maria DA, Sonohara S, Yamamoto M, Katayama MLH, Brentani MM. Increased expression of laminin receptors during myeloid differentiation. International Journal of Cancer. 1991;49(1):32–37.

43. Ihle JN. STATs: Signal Transducers and Activators of Transcription. Cell. 1996;84(3):331 – 334.

44. Gress TM, Kestler HA, Lausser L, Fiedler L, Sipos B, Michalski CW, et al. Differentiation of multiple types of pancreatico-biliary tumors by molecular analysis of clinical specimens. J Mol Med. 2012;90(4):457–464.

45. Rivlin N, Brosh R, Oren M, Rotter V. Mutations in the p53 Tumor Suppressor Gene Important Milestones at the Various Steps of Tumorigenesis. Genes & Cancer. 2011;2(4):466–474.

46. Cho Y, Gorina S, Jeffrey PD, Pavletich NP. Crystal structure of a p53 tumor suppressor-DNA complex: understanding tumorigenic mutations. Science. 1994;265(5170):346–355.

47. Fensterer H, Giehl K, Buchholz M, Ellenrieder V, Buck A, Kestler HA, et al. Expression profiling of the influence of RAS mutants on the TGFB1-induced phenotype of the pancreatic cancer cell line PANC-1. Genes, Chromosomes and Cancer. 2004;39(3):224–235.

48. Bhowmick NA, Neilson EG, Moses HL. Stromal fibroblasts in cancer initiation and progression. Nature. 2004;432(7015):332–337.

49. Kalluri R, Zeisberg M. Fibroblasts in cancer. Nature Reviews Cancer. 2006;6(5):392–401.

50. Ohuchida K, Mizumoto K, Murakami M, Qian LW, Sato N, Nagai E, et al. Radiation to stromal fibroblasts increases invasiveness of pancreatic cancer cells through tumor-stromal interactions. Cancer research. 2004;64(9):3215–3222.

51. Farrell TJ, Barbot DJ, Rosato FE. Pancreatic resection combined with intraoperative radiation therapy for pancreatic cancer. Annals of surgery. 1997;226(1):66.

52. Takada M, Nakamura Y, Koizumi T, Toyama H, Kamigaki T, Suzuki Y, et al. Suppression of human pancreatic carcinoma cell growth and invasion by epigallocatechin-3-gallate. Pancreas. 2002;25(1):45–48.

53. Albini A, Iwamoto Y, Kleinman HK, Martin GR, Aaronson SA, Kozlowski JM, et al. A rapid in vitro assay for quantitating the invasive potential of tumor cells. Cancer research. 1987;47(12):3239–3245.

54. Marchesi F, Monti P, Leone BE, Zerbi A, Vecchi A, Piemonti L, et al. Increased survival, proliferation, and migration in metastatic human pancreatic tumor cells expressing functional CXCR4. Cancer research. 2004;64(22):8420–8427.

